# Behavioral Analysis of Substrate Texture Preference in a Leech, *Helobdella austinensis*

**DOI:** 10.1101/409755

**Authors:** Rachel C. Kim, Dylan Le, Kenny Ma, Elizabeth A. C. Heath-Heckman, Nathan Whitehorn, William B. Kristan, David A. Weisblat

**Affiliations:** Department of Molecular and Cell Biology, 385 LSA, University of California, Berkeley, CA 94720-3200; Division of Biological Sciences, 3119 Pacific Hall, University of California San Diego, La Jolla, CA 92093; Present Address: Department of Ecology and Evolutionary Biology, University of California, Los Angeles, CA; Department of Physics and Astronomy, University of California, Los Angeles, CA

**Keywords:** *Helobdella*, leech, neuroethology, texture discrimination, touch-mediated behavior

## Abstract

Leeches in the wild are often found on smooth surfaces, such as vegetation, smooth rocks or human artifacts such as bottles and cans, thus exhibiting what appears to be a “substrate texture preference behavior”. Here, we have reproduced this behavior under controlled circumstances, by allowing leeches to step about freely on a range of silicon carbide sandpaper substrates. To begin to understand the neural mechanisms underlying this texture preference behavior, we have determined relevant parameters of leech behavior both on uniform substrates of varying textures, and in a behavior choice paradigm in which the leech is confronted with a choice between rougher and smoother substrate textures at each step. We tested two non-exclusive mechanisms which could produce substrate texture preference: 1) a Diffusion Trap mechanism, in which a leech is more likely to stop moving on a smooth surface than on a rough one, and; 2) an Anterior Choice mechanism, in which a leech is more likely to attach its front sucker (prerequisite for taking a step) to a smooth surface than to a rough one. We propose that both mechanisms contribute to the texture preference exhibited by leeches.

## INTRODUCTION

Although every cell will respond to mechanical stimuli at some intensity level (Belas 2014), animal nervous systems contain specialized mechanosensory neurons that mediate a tremendous diversity of behavioral responses to a wide range of mechanical stimuli (Kocer 2015). The relative simplicity of invertebrate systems makes them useful for identifying mechanosensory transducers and the behaviors they mediate. For example, studies in several leech species (*Hirudo medicinalis, H. verbana, Macrobdella decora, Haementeria ghilianii, Erpobdella obscura*) suggest that all leeches have three classes of segmentally iterated mechanosensory cells: three pairs of T cells, which respond to light touch; two pairs of P cells, which respond to moderate pressure, and two pairs of N cells, multimodal nociceptors that respond to acid, heat, and intense mechanical inputs such as pinching (Nicholls and Baylor 1968; Kramer and Kuwada 1983; Kuwada and Kramer 1983; Nusbaum and Kristan 1986; Pastor et al. 1996; Baltzley et al.2010). In addition to their different functional ranges of mechanical activation, T and P neurons differ in their dynamics: T cells respond best to transient or moving stimuli whereas P cells prefer sustained stimulation (Nicholls and Baylor 1968; Carlton and McVean 1995; Lewis and Kristan 1998b). Activation of T and P cells in different body locations (front, middle, and back) elicits different responses: shortening, local bending, and locomotion, both crawling and swimming (Kristan 1982; Kristan et al. 2005; Palmer et al. 2014).

At the behavioral level, the reflexive bending of the medicinal leech body wall in response to P cell stimulation has been used to reveal how location and intensity of mechanical stimuli are encoded by population coding vectors (Lewis and Kristan, 1998a 1998b). In addition, the initiation and modulation of shortening and swimming behaviors by mechanosensory inputs has been documented (Shaw and Kristan 1997; Kristan et al. 2005). Behavioral comparisons across leech species have revealed intriguing differences as well. For example, activation of P cells in the midbody of *H. verbana* produces a local bending response away from the site of stimulation whereas the same P cell activation in *E. obscura* causes them to roll themselves into a tight coil (Baltzley et al. 2010). In addition, activation of P or N cells in the back end of most leech species normally initiates locomotor behavior (crawling or swimming)(Kristan 1982; Palmer et al. 2014) but these responses are inhibited by feeding behavior specifically among various sanguivorous (blood-sucking) species, for which feeding opportunities are rare and of high energetic value, but not in predaceous species, for which feeding opportunities are more common and less valuable energetically (Gaudry et al. 2010).

Another leech behavior that would seem to be dependent on mechanosensation is suggested by the common knowledge (among leech gatherers at least) that submerged bottles, cans, and smooth rocks are excellent places to find leeches. At first glance, this behavior, which we term substrate texture selection, appears to involve a “preference” for smoother surfaces compared to rougher ones, but other explanations are possible. For example, it might be easier to detect and/or collect the animals from smooth objects. Moreover, even if the leeches do accumulate preferentially on smooth surfaces, this might involve a comparison of mechanosensory inputs by the animal or simply a “diffusion trap” in which leeches crawl longer or more rapidly on rough surfaces than on smooth surfaces. Here, we first demonstrate the existence of substrate texture preference under controlled conditions and then examine its behavioral basis, using a small glossiphoniid leech, *Helobdella austinensis* (Kutschera et al. 2013).

## MATERIALS AND METHODS

### Animals

Adult leeches (*Helobdella austinensis*; Kutschera et al., 2013) were obtained from a laboratory breeding colony kept at room temperature (ca. 21-25° C) in artificial pond water (1% artificial seawater [Salinity for Reefs; AquaVitro]. Animals are housed in freely breeding groups of mixed ages, ranging from a few dozen to a few hundred individuals in 0.5-2 liter glass or plastic containers. Animals were fed 2-5 times per week by adding several grams of frozen midge larvae (BloodWorms; Hikari or Omega One) to each bowl and then changing the water after the animals had fed to satiety (2-5 hours). The state of satiety is readily judged by the extent to which their guts--easily visualized through the translucent body wall--are filled with red pigment from the ingested midge larvae.

### Split Dish Substrate Texture Preference Assay

Leeches were given at least 2 hours to feed before each trial. After feeding, individuals to be used were blotted dry and weighed using an analytical scale. Animals bearing embryos or juveniles were not used.

Assay chambers consisted of circular plastic petri dishes (Falcon; 60mm diameter, 15mm depth), the internal surfaces (bottom and sides) of which were lined with waterproof silicon carbide sandpaper (3M Wetordry^™^), which was glued to the plastic surface by either vinyl or spray contact adhesive (3M). The two halves of each chamber were lined with different grades of sandpaper, ranging from 80 to 1500 grit, respectively. Each dish was filled with 13mL of spring water to a depth of 5mm and placed inside a light-proof container illuminated from above with infrared (IR) LEDs. Between each trial, dishes were rinsed thoroughly with deionized water. For each assay, one leech was placed in the center of the rougher grade sandpaper at the beginning of the trial. Each trial lasted 25 minutes, and started once the light-tight container was placed over the dish. Trials in which an individual failed to approach the smoother substrate were discarded. A new leech was used for each trial. All trials were carried out at room temperature (RT; roughly 22°C).

Images were acquired using a D-Link Wi-Fi camera (1 MPixel) or a Samsung Galaxy S4 Zoom camera (16 Mpixels; modified by removing the factory-installed IR filter) supported inside the light-proof container, above the dish and level with the IR LED illumination. Pictures were taken once every 10 seconds for 25 minutes (150 frames per trial).

Data analysis. For each trial, the time spent on each side was calculated, with the position at each time point defined by the location of the posterior sucker. Because pictures were taken once every ten seconds, which is short relative to the average step frequency, it was assumed that there was at most one step taken within the ten seconds following each picture. Percentage of time spent on each grade was calculated, and deviation from random preference (50% of time spent on each grade) was assessed by a t-test with a post-hoc Bonferroni correction for multiple comparisons.

### Checkerboard Substrate Texture Preference Assay

Leeches were prepared and selected according to the protocol described above. Assay chambers consisted of square plastic petri dishes (Fisherbrand; 100mm x 100mm), the dishes were lined with waterproof silicon carbide sandpaper (3M Wetordry^™^), as described above, except that the bottoms of the dishes were lined with a 6×6 checkerboard pattern with alternating squares (13mm x 13mm) of two selected textures. In each checkerboard chamber, the sides were lined with the rougher of the two textures under comparison, to discourage the leech from attaching to the sides of the chamber. Each dish was filled with 35 mL of spring water to a depth of 5mm, and placed in the same light-proof container as above. Between each trial, chambers were rinsed thoroughly with deionized water. Assays began with one leech placed on a center rough or smooth square under the light-tight container, and ended after 5 minutes. A new leech was used for each trial. All trials were carried out at RT, roughly 22°C. Videos were acquired according to the same protocol as above.

#### Data analysis

For each trial, each step taken was sorted into one of four categories, based on the substrate texture at the origin and the termination of the step: from smooth to smooth, from smooth to rough, from rough to rough and from rough to smooth--this corresponds to four categories of transitions, abbreviated SS, SR, RR and RS, respectively. For each experiment, we evaluated the preferences of the leeches for each of the four possible transitions by comparing the total number of transitions in each category to the numbers predicted by a Monte Carlo simulation modeling the behavior of an identical number of simulated leeches, with the same starting conditions (i.e., starting on the rough or smooth surface), and the same number of transitions (steps) for each leech. In our Monte Carlo simulations, we assumed a memoryless process with fixed, time-independent probabilities for each type of transition: p(RS), p(RR) = 1-p(RS), p(SR), and p(SS) = 1-p(SR).

We then found the values of these probabilities that best match the data. For each experiment, we define “best match” by an objective function equal to the sum of the squares of the differences in each of the four total transition counts between a simulation with a given set of transition probabilities and the observed data; in the limit of Gaussian errors, this provides an optimal maximum-likelihood estimator (Neyman and Pearson, 1933). We estimated these expected values by taking the means of the transition counts from 50 simulations run for each set of transition probabilities in a grid with test points corresponding to differences in probability of 0.01.

To calculate *p*-values, we first ran simulations of the parameters p(RS) = p(SR) = 0.5, corresponding to the leeches moving randomly between the two substrate textures. For each simulation, we tabulated the values of the objective function tested against the p(RS) = p(SR) = 0.5 hypothesis, which generates the statistical distribution of how well the behavioral data matched expectations by chance. We then counted the fraction of these simulation runs that, by chance, fit p(RS) = p(SR) = 0.5 as well as or more poorly than the data to give the final quoted *p* values.

### Uniform Substrate Texture Response Analysis

Leeches to be used in these studies were anesthetized by immersion in cold water (5°C) to allow for removal of any attached eggs and juvenile leeches, then maintained individually in 9×50mm plastic petri dishes filled with artificial pond water for at least 24 hours at room temperature before testing. Animals were fed on bloodworms (Omega One) 24 hours prior to experimentation.

#### Step frequency assay

Assay chambers (Falcon 9×50mm plastic petri dishes) were constructed as described above except that their insides were covered with a single grade of silicon carbide (3M Wetordry^™^) sandpaper (80, 150, 320, 400, 600, or 1500 grit); unlined petri dishes served as controls. Each leech was placed in a pond water-filled chamber and observed for 30 minutes after its first complete step. Experiments were initially conducted in red light (Fostec 8375 light source through Bleyer red gift wrapping paper), because leeches are insensitive to red light (Jellies 2014). Some of the same leeches were also tested in white light, and their stepping behavior on different sandpaper grits was statistically indistinguishable from those tested in red light. Thus, for convenience, later trials were run exclusively in white light. Each leech was tested on every grade of sandpaper, in a random order, then kept for at least 30 minutes in an unlined petri dish before being transferred onto another grade of sandpaper for the next testing session.

#### Head withdrawal assay

Using the same chambers, we counted the number of times the animal lifted its anterior sucker from the surface of the sandpaper before re-attaching this sucker to complete a step. Viewed from above (i.e., the dorsal surface of the leech), head raises were distinct from other movements of the anterior because the ventrally-located anterior sucker became visible. Head raises accompanied by rapid shortening of the body were classified as head withdrawals. These experiments were conducted exclusively in red light, and animals were given 30 minutes of recovery time in plastic petri dishes before being placed into another sandpaper chamber. Animals were recorded using a Point Grey Flea3 USB camera mounted on the trinocular head of a dissecting microscope (15 fps).

## RESULTS

### Heavy but not light *H. austinensis* accumulate preferentially on smoother surfaces

The fact that, in the wild, leeches are often found on the undersides of submerged objects such as cans, bottles, vegetation, or smooth rocks led us to ask if leeches have the ability to distinguish between different textures, as judged by the ability to reliably exhibit a preference for one texture over another under controlled conditions.

In contrast to *Hirudo*, leeches in the genus *Helobdella* neither swim nor exhibit the peristaltic crawling movements used by earthworms. Instead, they locomote by an inchworm-like stepping (Stern-Tomlinson et al. 1986). Each definitive step is often preceded by a side-to-side *scanning behavior* of the anterior end, as if it were sampling the surrounding area before taking a step. The European medicinal leech, *Hirudo verbana,* produces similar scanning behavior (Harley and Wagenaar 2014).

To determine if leeches exhibit a substrate texture preference, we placed individual *H. austinensis* leeches of various sizes in dishes lined with two grades of sandpaper (Fig. 1A,), and observed their positions at ten-second intervals for 25 minutes; data from one trial for one leech is shown in Fig. 1B. This animal took 13 steps in the 25-minute cycle. The radius of each blue circle is proportional to the length of time that the back sucker occupied that location, and thus represents the time between steps. The large blue circle at the 13^th^ step indicates that the animal stayed in this location for several minutes until the end of the trial. The smaller green circles represent the locations of the front sucker. Their small size and scattered locations indicate the relatively high frequency of the scanning movements. From many such trials, we calculated the percentage of time spent on each texture, by comparing the sum of the areas of the blue circles on each of the substrates. To eliminate the possible influence of uneven lighting, these experiments were conducted in darkened chambers using IR illumination (to which leeches are insensitive [Jellies 2014]) and an IR-sensitive camera to record the animals’ movements. The trial duration of 25 minutes was selected as an interval after which most animals had ceased moving following the arousal caused by the initial handling.

**Figure 1.**
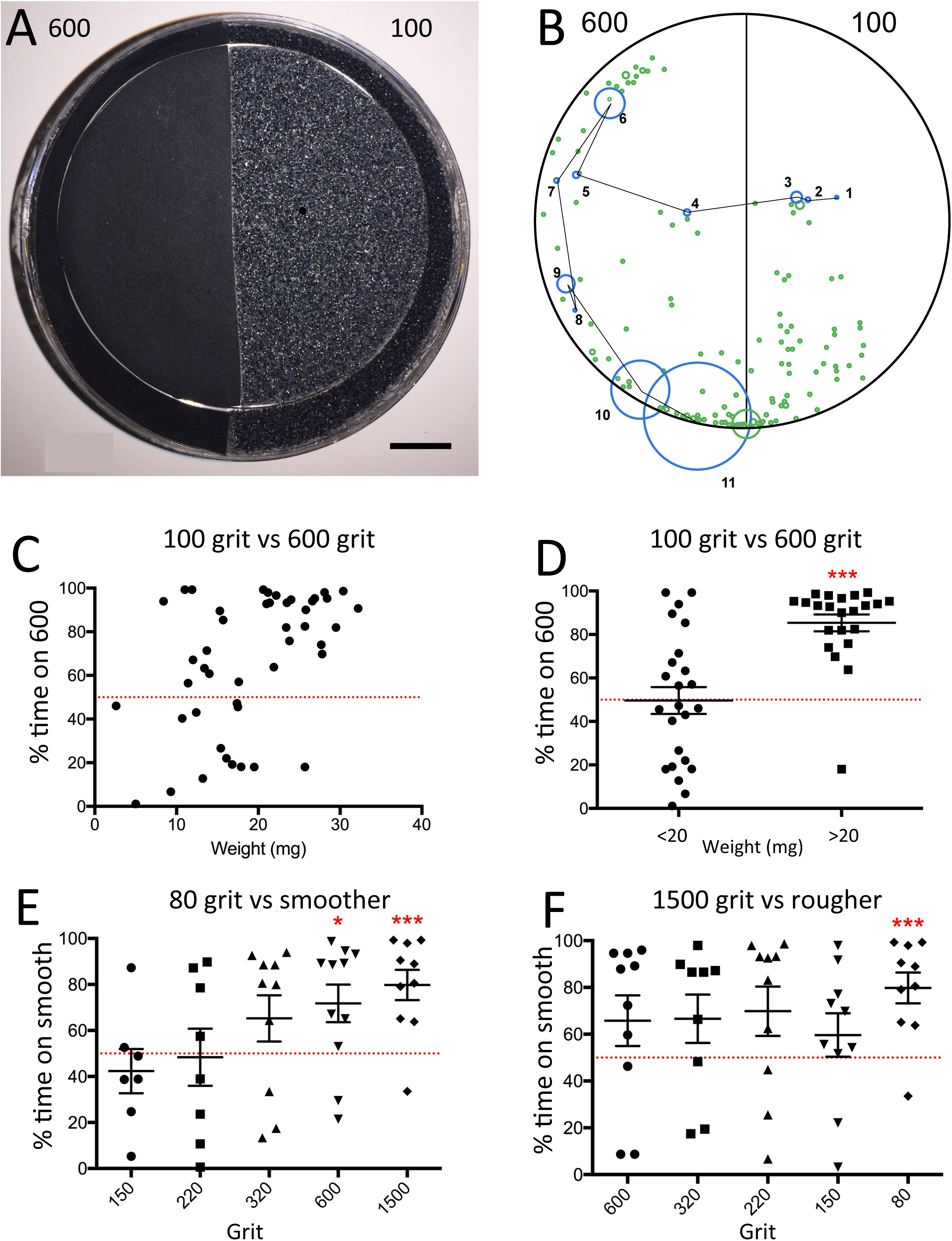
Heavier leeches end up preferentially on smoother substrates. **(A)** Representative texture preference assay chamber, photographed from above, with 600 grit (left) and 100 grit (right); scale bar, 1 cm. **(B)** Raw data of leech sucker placement during a trial of 25 minutes. Blue circles indicate posterior sucker placement, and green circles indicate anterior sucker placement. Circles increase in radius proportionally to time spent in one position. **(C, D)** Plotted results from 46 trials in which individual leeches were placed on the rougher (100 grit) surface and allowed to navigate the apparatus freely in the dark (IR illumination) for 25 min; each point represents a single trial. In **(C)**, the percent of the trial durations spent on the smoother substrate are plotted as a function of the weights of the 48 animals tested. In **(D)**, the results for animals weighing less than or greater than 20 mg are grouped separately. Leeches weighing <20 mg spent as much time on the rough and smooth sandpapers, whereas all but one of the leeches over 20 mg spent more than 50% of time on the smoother sandpaper (*p* < 0.001). **(E, F)** Plotted results of a grand total of 85 similar experiments in which leeches (all greater than 20 mg) experienced 80 grit substrate texture on one side of the chamber and a range of smoother substrates (150, 220, 320 600 and 1500 grit) on the other **(E)**, or 1500 grit substrate texture on one side and a range of rougher substrates (600, 320 220, 150 and 80) on the other **(F**; data for the 80 vs 1500 grit comparison are included in both plots). In this experimental paradigm, only leeches confronting the more dramatic differences (80 vs 1500 and 150 vs 600) exhibited a statistically significant preference for smooth surfaces, but there is an apparent trend for preferring the smoother surface in most comparisons. Most of the outlying data points represent animals that failed to cross to the smoother side until late in the trial.

In the first series of experiments, we tested leeches on a fixed comparison of 600 grit (finer) versus 100 grit (coarser) sandpaper. These experiments revealed a weak preference for the finer grit (Fig. 1C). Closer analysis revealed that lighter leeches (in the weight range of 2-20 mg) showed no clear preferences, but that the heavier leeches (20-35 mg) exhibited a significant preference for the smoother substrate (*p* < 0.001) (Fig. 1D). One possible explanation for this preference is that smaller, lighter individuals do not experience a force from the substrate that is sufficient to activate appropriate mechanoreceptors, whereas larger, heavier individuals do. Alternatively, it could be that the surface area of the sucker somehow impacts the animal’s perception of the substrate texture or its ability to establish a firm grip. Whatever the cause, based on their more reliable substrate texture response, only leeches weighing more than 20 mg were used for further experiments.

### Leeches may discriminate across a range of substrate textures

To assess the leeches’ ability to discriminate between different substrate textures, we used the same split dish assay to test for substrate preferences in pairwise comparisons of various textures ranging from coarse (80 grit) to fine (1500 grit). In these experiments we compared each extreme of the spectrum against all other values, i.e., 80 vs 150, 220, 320, 600, and 1500 grit, as well as 1500 vs 150, 220, 320, and 600 grit. As expected, the smoother surface was favored in all assays that revealed a significant substrate texture preference, but the ability of the leeches to discriminate textures was limited (Fig. 1E). In this assay, the leeches did not exhibit a statistically significant discrimination between 80 grit and 150, 220 or even 320 grit. Leeches did spend significantly more time (an average of 71.79%) on the 600 grit than on the 80 grit sandpaper (*p* < 0.05), and even more time on the 1500 grit (an average of 79.81% *p* < 0.001) rather than the 80 grit paper. Conversely, in experiments where the 1500 grit surface was compared against all other surfaces, the leeches preferred the smoother surface on average, but the preference was statistically significant only for 1500 vs 80 grit (Fig. 1F).

These results demonstrate that leeches do prefer smoother surfaces, at least when there is a large difference in the texture of the substrate. To test how leeches end up on the smoother surface in the bisected smooth/rough petri dish, we considered two general classes of behavioral possibilities: (1) Diffusion Trap, in which the leech steps in random directions (“diffusion”), and then accumulates preferentially on the smoother surface (the “trap”), and; (2) Smoothness Selection, in which the leech preferentially moves onto the smoother surface when confronted with the rough/smooth border. These two possibilities are not mutually exclusive. We tested for the first strategy by monitoring leeches’ step frequencies on a surface of uniform roughness so that their only choice was whether to continue stepping or to stop. We tested for the second possibility by providing walking leeches with so many rough/smooth borders that they faced a choice with every step.

### Increased step frequency, and duration of locomotory episodes, on rougher surfaces is consistent with a Diffusion Trap mechanism

We addressed the “diffusion trap” behavioral strategy, in which leeches slow down or stop on smooth substrates, by quantifying the stepping behavior of individual leeches on uniform substrates across a range of textures from 80 grit to 1500 grit. Specifically, we measured the step rate (number of steps per each 2-minute interval [Fig. 2A]) by individual leeches placed into unlined plastic petri dishes (representing a smooth control surface) and in dishes lined with either 80, 150, 320, 400, 600, or 1500 grit sandpaper. Despite the fact that leeches exhibit negative responses to short-wavelength light (Jellies 2014; Bisson 2011; Harley et al 2011), step frequency was not significantly affected by the color (red vs. white) of the light in which animals were tested (data not shown). During the first 8 minutes, step frequencies decreased slightly on all substrate textures. But while the step frequencies for smoother substrates continued to decline, those measured on the two roughest substrates remained steady (150 grit) or even increased with time (80 grit). Thus by 16 minutes, the step frequencies for animals on 80 and 150 grit were clearly different from each other and from those on smoother substrates.

**Figure 2.**
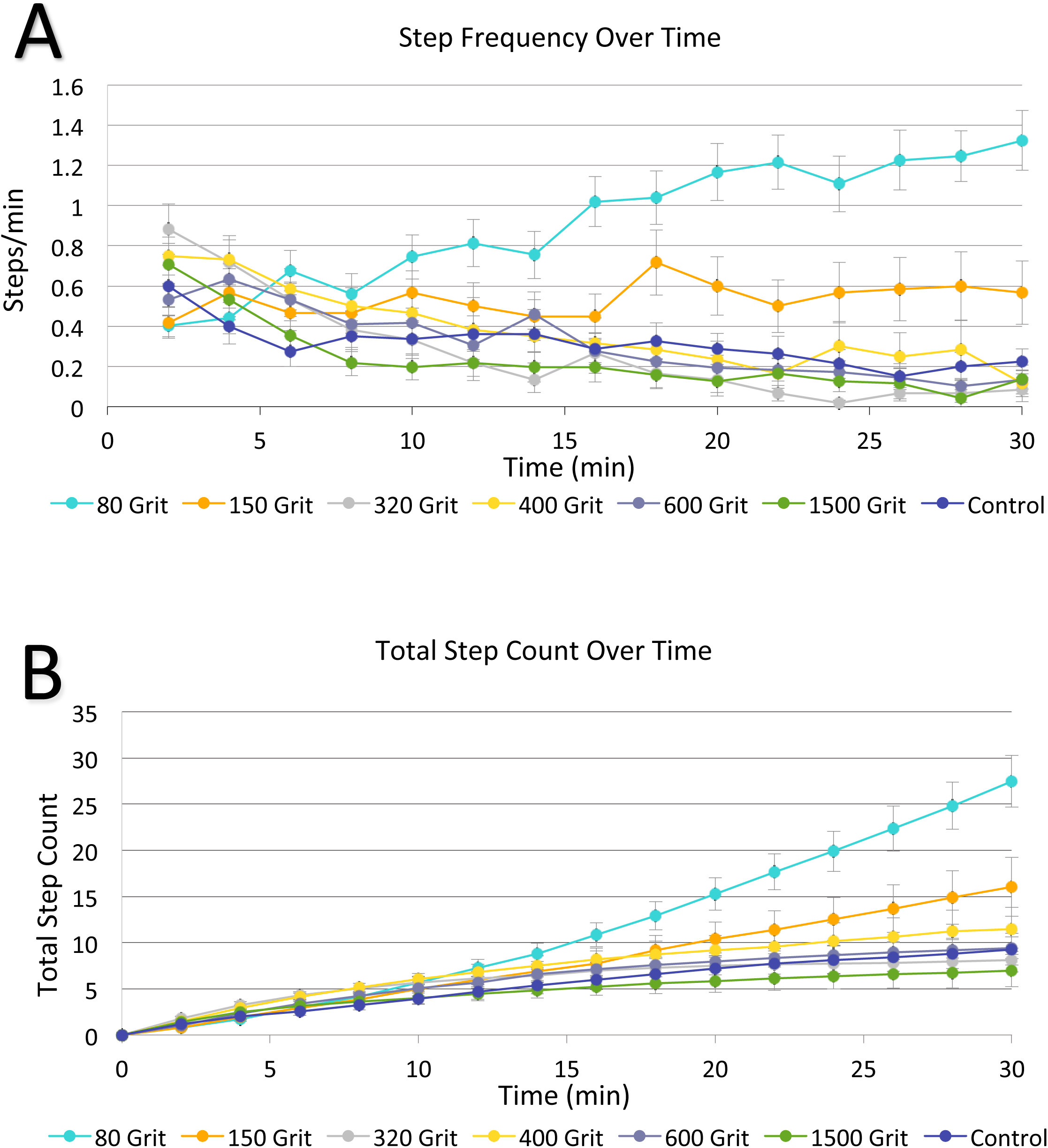
Step frequency and duration of locomotory episodes increased on rougher substrates. **(A)** Average number of steps per minute during 30 minute trials on uniform substrates covering a range of textures (80, 150, 320, 400, 600 and 1500 grit sandpaper, or a control smooth surface with no sandpaper). While step frequencies on the four finest grits are indistinguishable from control, decreasing over time, step frequencies remain roughly constant on 150 grit and increase over time on 80 grit (the coarsest substrate). **(B)** The same data as in A plotted as cumulative steps over time. The total number of steps taken over the 30 minute trial on 80 grit was significantly higher than all other conditions tested (*p* < 0.01 vs 150 grit, *p* < 0.001 for all other grits, T-Test). The step totals on 150 grit sandpaper also deviated from several other conditions (*p* < 0.05 vs 320 and 1500 grit sandpaper conditions, T-Test). Roughly half the trials were conducted in white light and half in red light, to which leeches are not sensitive (Jellies 2014). Because there were no statistical differences between these two conditions, the data were combined. Each data point represents the average of between 29 and 51 trials.

For a more rigorous statistical analysis of these data, we plotted the cumulative number of steps over the 30-minute test period (Fig. 2B). Consistent with the effects of grit roughness on step frequency (Fig. 2A), the total number of steps taken by animals on 80 grit sandpaper (Fig. 2B) deviates significantly from those taken on all other surfaces studied (*p* < 0.01 vs 150 grit, *p* < 0.001 for all other grits, Student’s T-Test). The number of steps on 150 grit sandpaper also differs significantly from other conditions (*p* < 0.05 when compared against 320 and 1500 grit conditions, T-Test). Interestingly, the differences in step number are not apparent immediately. The cumulative step total on 80 grit sandpaper begins to diverge from 150 grit after 22 minutes, from 400 grit paper after 18 minutes, from 320 and 600 grit after 16 minutes, and from from both 1500 grit and control after 12 minutes (*p* < 0.05, all comparisons). There are more total steps on 150 grit sandpaper beginning at 20 minutes when compared to 1500 grit sandpaper and after 26 minutes when compared against 320 grit sandpaper (*p* < 0.05).

The behavior of leeches on uniformly rough surfaces is consistent with the Diffusion Trap mechanism: leeches exhibit higher step frequency (Fig. 2A) and longer stepping episodes (Fig. 2B) on rougher surfaces and tend to settle on smoother surfaces.

### A checkerboard assay provides evidence for Smoothness Selection

To generate an experimental environment in which individuals faced a choice between rough and smooth substrates on every step, we placed leeches on checkerboard substrates constructed of alternating rough and smooth squares, the size of which (13 by 13 mm) was chosen to approximate the length of the largest resting leech (Fig. 3A; see Materials and Methods for details).

**Figure 3.**
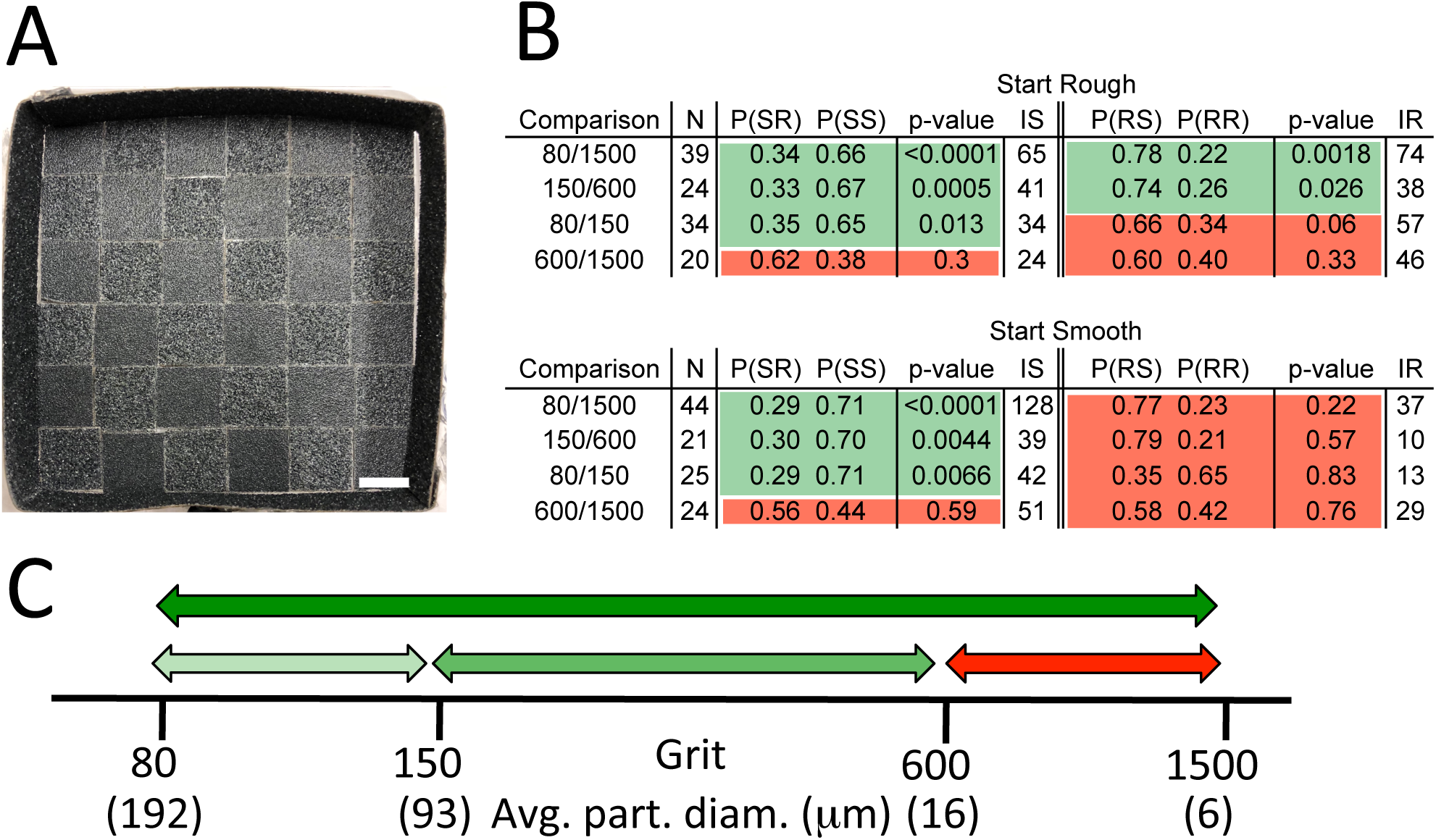
A checkerboard assay reveals that leeches can discriminate 80 vs 1500 grit, 80 vs 150 grit, and 150 vs 600 grit substrates. **(A)** Representative checkerboard assay apparatus, showing alternating rough and smooth squares, viewed from above; scale bar, 1 cm. (Supplemental video 1 shows leeches navigating the apparatus.) **(B)** Tabular summary of checkerboard experiments. For each comparison (e.g. 80 grit vs 1500 grit squares), two series of trials were conducted, started by placing a leech on a rough square (upper table) and on a smooth square (lower table), respectively. In each row of the table, N is the number of usable trials; IS and IR are the total number of steps initiated from smooth or rough squares in those trials, respectively. From all the steps taken in each set of trials, we used a Monte Carlo simulation approach to estimate the probabilities of the four possible transitions, i.e., from smooth to rough, from smooth to smooth, from rough to smooth and from rough to rough--p(SR), p(SS), p(RS) and p(RR)--and to test whether the data were consistent with leeches making the four transitions randomly. As an aid to the reader, findings with p-value < 0.05 are highlighted in green, and those with p-value >0.05 are highlighted in red (see Materials and Methods for details). **(C)** Qualitative summary of the results, showing that the leeches were able to distinguish between rough and smooth surfaces in all comparisons (green arrows) except 600 grit vs 1500 grit (red arrow) and that larger differences led to stronger preferences (different shades of green).

First, we conducted five-minute trials with 100 different leeches individually on a checkerboard of the roughest and smoothest textures (80 grit and 1500 grit), starting 50 animals on a rough square and the other 50 on a smooth square. Sixteen of these 100 trials yielded no data either because of technical problems or because the leech took no steps during that trial. In the remaining 84 trials, casual inspection revealed that the leeches made more steps onto smooth squares than onto rough ones (e.g., Supplemental Video 1). Given the apparent ability of the leeches to discriminate the 80 vs 1500 grit squares, we used similarly patterned substrates to test their ability to discriminate rough from smooth textures on three intermediate pairs of substrate textures: 80 vs 150 grit, 150 vs 600 grit and 600 vs 1500 grit.

To quantify leech stepping behavior in the checkerboard assays, we sorted the steps from each experiment into four separate categories (transitions) according to the substrate texture at the origin and destination of each step (Summaries of raw data provided in Supplement). Thus, the four possible transitions were rough-to-rough (RR), rough-to-smooth (RS), smooth-to-rough (SR), and smooth-to-smooth (SS). For each experiment, we evaluated the preferences of the leeches for each of the four possible transitions by comparing the total number of steps in each category to the numbers predicted by Monte Carlo simulation with a given set of transition probabilities p(RR), p(RS), p(SR) and p(SS), and then varied the transition probabilities to get the best fit to the observed experimental data. Statistical significance (p-value) of the results was measured by comparing the data to a model in which each transition was equally likely, i.e., p(RR) = p(RS) = p(SR) = p(SS) = 0.5 (see Materials and Methods for details).

As expected from casual observations (Supplemental Video 1), leeches readily discriminated between 80 grit and 1500 grit (Fig.3B). The statistical analysis of the behavior was constrained by the fact that, for trials started by placing animals on smooth squares, the number of steps initiated from rough squares (Fig. 3B, Start Smooth,IR) was much smaller than the number of steps initiated from smooth squares (Fig. 3B, Start Smooth, IS), because of the leeches’ tendency to avoid the rough surface. Note that the Start Smooth trials comparing 80 vs 1500 grit and 150 vs 600 grit averaged less than one step initiated from a rough square (IR) per trial (Fig. 3B). Apart from this complication, however, the differences between the observed behaviors and those predicted from a model with equally likely transitions was highly significant (Fig. 3C).

Leeches also discriminated between rough and smooth surfaces in two of the three other comparisons as well, namely, 80 vs 150 grit and 150 vs 600 grit (Fig. 3B, C). The strength of these behavioral discriminations was less pronounced than for the 80 vs 1500 comparison (as judged by their generally higher p-values), but still statistically significant for the trials with sufficient numbers of steps. In contrast, for the third comparison (600 vs 1500 grit), the step distributions generated by the best fit Monte Carlo simulations were not significantly different than those predicted by transition probabilities of 0.5 under any condition, indicating that the leeches did not discriminate between 600 and 1500 grit squares (Fig. 3B, C).

We also considered the possibility that differences in step duration might influence the behavioral outcomes in the checkerboard experiments. For example, if leeches made quicker steps from rough squares to smooth squares (RS) than from smooth squares to rough squares (SR), they would spend more time on smooth surfaces than on rough surfaces. To address this possibility we measured step duration, defined as the interval between releasing the rear sucker in successive steps. Thus, the step duration included both scanning behavior between rear sucker releases, as well as other behaviors, and complete immobility. Even for the most extreme (80 vs 1500) comparison, we found no significant difference between the average duration of RS steps (35 +/- 62 sec; n = 76) and SR steps (45 +/- 54 sec; n = 51).

### Frequency of head withdrawal during scanning correlates with substrate texture

A single crawl step by a leech consists of six distinct components: (1) attaching the rear sucker to the substrate near the attached front sucker, (2) release of the front sucker, (3) extending the whole body, (4) attaching the front sucker, (5) releasing the back sucker, (6) shortening the whole body to bring the rear sucker near the front sucker again. Sometimes these steps are repeated in direct succession to produce a rapid crawling behavior (Stern-Tomlinson et al. 1986). More often, however, components 3 and 4 are separated by a much more variable, elective component called “scanning”. Scanning behavior consists primarily of back-and-forth sweeping motions of the front sucker and anterior midbody (Fig. 4A; Supplemental video 2), during which the ventral surface of the anterior sucker makes frequent contact with the substrate. Scanning also includes occasional “head raises”, in which the animal explores above the plane (e.g., starting at roughly 4 seconds and 19 seconds in Supplemental video 2); less frequently, the scanning behavior is punctuated by rapid and pronounced retractions that lift the anterior end away from the surface and simultaneously shorten the body (Figure 4B, C; e.g., starting at roughly 9 seconds in Supplemental video 2), a behavior that we call “head withdrawal”. The speed and vigor of the head withdrawal movements gives the impression that the animal is recoiling from a noxious stimulus.

Our impression was that leeches make more head withdrawals on rough surfaces than on smooth ones. We tested this possibility by monitoring leeches crawling on uniform substrates of differing roughness. Specifically, we placed individual leeches onto the bottoms of plastic petri dishes covered with 80, 150, 320, 400, 600, or 1500 grit sandpaper, or onto plastic alone as a control, smooth surface. We counted the number of head withdrawals made by 12 animals as they took 5 steps on each surface. We found that the number of head withdrawals was greatest for the two coarsest sandpapers and diminished on smoother sandpapers (Fig. 4D). During head withdrawals, a leech cannot attach its front sucker to produce a step. Thus, the higher number of head withdrawals on the coarse surfaces could result in a higher probability of the anterior sucker attaching to smoother surfaces when it has access to substrates that differ in texture.

**Figure 4.**
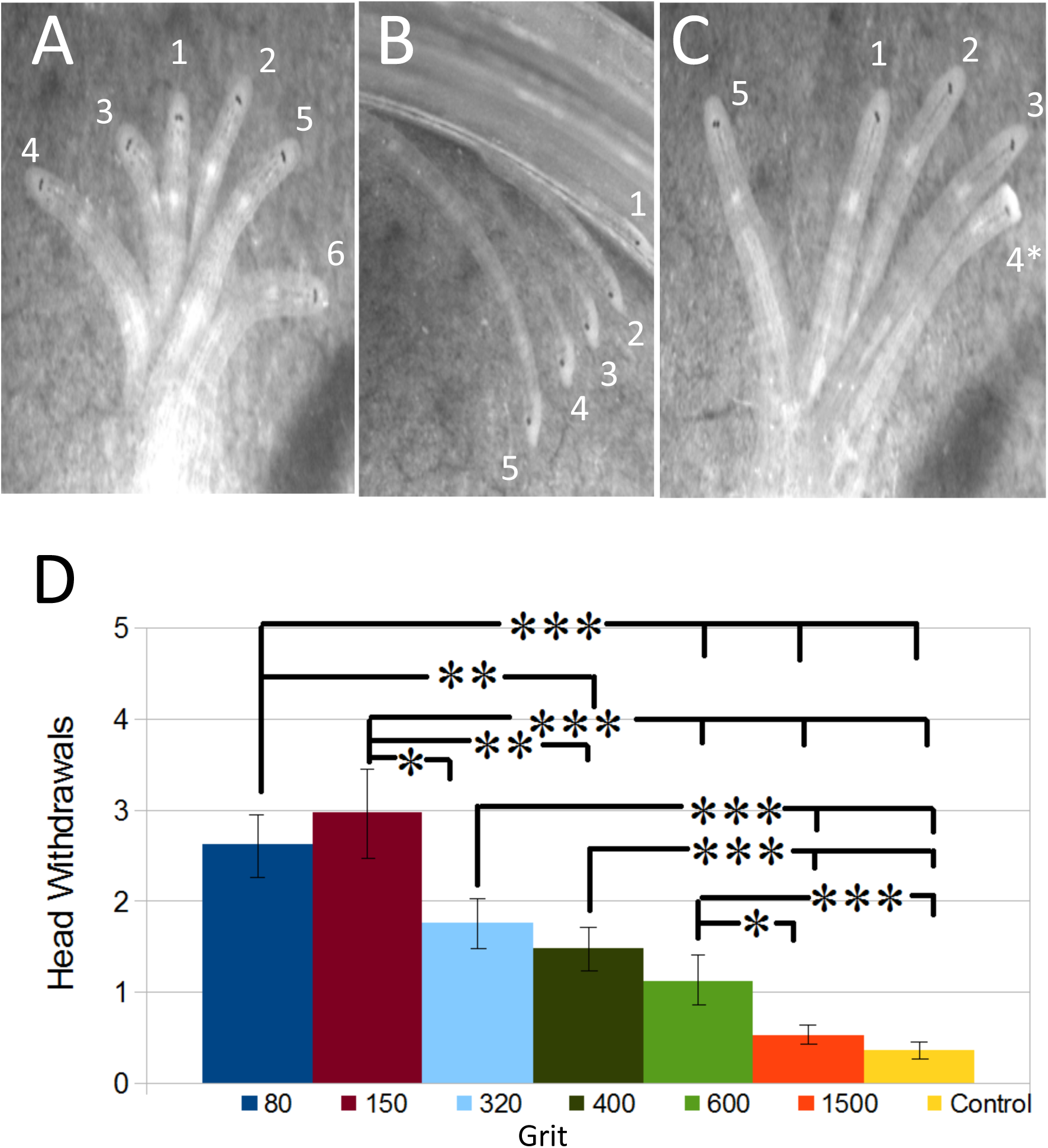
“Scanning”, an optional exploratory component of stepping behavior, includes “sweeping”, “head raising”, and “head withdrawal” behaviors; head withdrawals are more frequent on rougher substrates. **(A-C)** Images constructed by overlaying individual digital images taken at 5 or 6 successive time points, in the sequence indicated by the numbers associated with the images, separated by variable time intervals. (**A)** Sweeping behavior, viewed from above. The time intervals between positions 1-2, 2-3, 3-4, 4-5, and 5-6 were 2s, 4s, 10s, 4s, and 4s, respectively. (**B**) A single episode of head raising behavior, viewed from the side, by an animal attached to the side of the petri dish. Frames 1-5 represent successive 400 ms time points. **(C)** Scanning behavior, viewed from above, with a head withdrawal (asterisk). Time intervals between positions 1-2, 2-3, 3-4, 4-5 are 1s, 1s, 1s, 11s, respectively. **(D)** Plot shows the number of interstep head withdrawals during 5 steps by each of 12 animals on each of 6 substrate textures and a smoothness control (no sandpaper; n = 60 steps per condition). The interstep head withdrawal frequency on 80 grit differed significantly from those on 400, 600 and 1500 grits and the control (*p* < 0.01 for 80 vs 400 grit, *p* < 0.001 for 80 vs 600, 1500, and control). Head withdrawal frequency on 150 grit significantly differed from those on all smoother conditions (*p* < 0.05 for 150 vs 320; *p* < 0.01 for 150 vs 400; and *p* < 0.001 for 150 vs 600, 1500, and control). Both 320 and 400 grit conditions were found to be significantly different from the 1500 grit and control conditions (*p* < 0.001 for both). Leeches raised their heads more frequently on 600 grit sandpaper than on either 1500 grit sandpaper (*p* < 0.05) or the control surface (*p* < 0.001). On the diagram, * = *p* < 0.05, ** = *p* < 0.01, and *** = *p* < 0.001.

## DISCUSSION

Our behavioral studies started with the observation that leeches tend to settle onto smooth surfaces, such as smooth rocks or stems of reeds, as well as on artifacts such as bottles and cans. In the first set of experiments (Fig. 1), we established that we could produce this behavior in a controlled setting, by allowing leeches to move about on a substrate divided into smoother and rougher halves using different grits of sandpaper. These studies showed that the leeches moved around the fairly large chamber and, if the difference in roughness was sufficiently great, tended to settle on the smoother side.

### Basis for settling on smooth surfaces

Our behavioral data showed that leeches have two distinct mechanisms for settling on smooth surfaces: they move more slowly on a smooth surface than on a rough one (Fig. 2), a mechanism we call Diffusion Trap; and they select patches of smoother surface when presented with a choice of smooth and rough substrates (Figs. 3), which we call Smoothness Selection. We considered three possible mechanisms that could underlie Smoothness Selection: (1) Spatial Comparison, in which the leech compares the textures felt at different body locations, most likely at its anterior and posterior suckers because these are the only body parts that are in almost continuous contact with the substrate; (2) Anterior Choice, an increased likelihood for a leech to attach its anterior sucker to smoother surfaces during its scanning behavior before each step; (3) Temporal Comparison, in which a leech compares the texture that it is experiencing right now with the texture that it experienced at some previous time. Note that these possibilities are not mutually exclusive. Our experiments were aimed at testing these possibilities.

#### Spatial Comparison and Anterior Choice mechanisms

The experiments on the rough/smooth checkerboards (Fig. 3) address the Spatial Comparison and Anterior Choice possibilities. In effect, the checkerboard provides an ongoing choice between rough and smooth because the leech can reach squares with either texture at every step. Supplemental Video 1 shows that the crawling leech readily reaches the 8 adjacent squares, and in no case did a leech take a step onto the square to which the back sucker was attached. In both mechanisms, the anterior sucker samples possible substrates, but a Spatial Comparison mechanism evaluates input from two locations (presumably the front and back suckers), whereas an Anterior Choice mechanism uses input from only the front sucker. In fact, the probability of stepping onto a smooth surface is similar whether the back sucker is on a smooth surface (a SS step) or a rough one (an RS step) (Fig. 3B). These data support the Anterior Choice mechanism as a contributor to settling on a smooth surface.

#### Temporal Comparison mechanism

The Anterior Choice mechanism could use an absolute measure of sensory activation or it could calculate the difference in sensory activation at two different times. Such a difference might be measured at different times within a single scan or by measuring the average activation in one scan compared to a second scan. Our one result that bears upon this question is that we found no significant difference between the average duration of steps from rough to smooth squares (35 +/- 62 sec; n = 75) and those from smooth to rough squares (45 +/- 54 sec; n = 51), despite a large difference in the probabilities of RS and SR steps. Given the large variability, this is not a definitive test of the Temporal Comparison model, but is would be surprising if two such different decisions took the same amount of time. Hence, our data are weakly against Temporal Comparison as a basis for deciding where to step.

#### Relative importance of Diffusion Trap and Anterior Choice mechanisms

The fact that leeches continue to move on a rough surface for much longer times than on smoother surfaces (Fig. 2A) is consistent with the Diffusion Trap mechanism, which would increase the likelihood that a leech would settle on a smooth surface rather than a rough one. The difference between the number of steps on the coarsest and smoothest grits is highly significant--about 20 steps after 30 minutes (Fig. 2B). Each step is nearly a full extended body length, so a 20-step difference could move the leech into very different territory. The Anterior Choice mechanism would direct these additional steps into smoother substrates. Thus, these two mechanisms, working together, would provide both an impetus to start and stop crawling (Diffusion Trap) and an directionality that could lead to a more desirable, smoother substrate (Anterior Choice).

### Possible implementation of Anterior Choice and Diffusion Trap mechanisms

The Anterior Choice model implies that leeches somehow react to the roughness of a surface using their anterior suckers, the most motile part of the leech, and the Diffusion Trap mechanism suggests that the sensitivity of crawling behavior to tactile receptors in the anterior suckers does not wane, and may even increase, over time (Fig. 2). The characterization of the behavior of the front end of these leeches shows that they often use left-to-right scanning movements along the substrate before taking a step, and they also make occasional head withdrawals, which lift their head off the surface of the substrate (Fig. 4A; Supplemental video 2). The fact that these head withdrawals are more frequent on a rough substrate (Fig. 4B) suggests that these movements are a component of the Anterior Choice mechanism.

The mechanosensory system of the leech provides some possibilities about how these mechanisms might be implemented. (NB: Most of this information has been garnered from work on other species than *Helobdella*, primarily European medicinal leeches, but mechanosensory neurons have been recorded in many other leech species (Kramer and Kuwada 1983; Kuwada and Kramer 1983; Lent 1985; Nusbaum and Kristan 1986; Baltzley et al. 2010) without significant differences among the species.) In each of the leech’s 21 midbody segments there are 3 pairs of T cells (respond to very light touch), 2 pairs of P cells (respond to pressure on the skin), and 2 pairs of N cells (respond to strong mechanical, chemical, and thermal stimuli) (Nicholls and Baylor 1968; Carlton and McVean 1995). The same neurons are present in the head brain, which innervates the anterior sucker (Yau 1976). In general, behavioral responses require stimuli sufficient to activate P cells, with a boost from the activation of T cells; activation of N cells produces qualitatively different responses (Kristan 1982; Palmer et al. 2014). A reasonable hypothesis, therefore, is that the T and P cells are being activated as the anterior sucker is scanned over the substrate, and that occasional activation of N cells by the sharp edges of the grit could trigger the rapid head withdrawals.

Why, in fact, does a leech crawl at all? In nature, among other reasons, they crawl seeking food (Sawyer 1981; Lent 1985; Harley et al. 2013, 2014), finding a mate (Sawyer 1981), interacting with other leeches (Bisson and Torre 2011), and when agitated (Willard 1981). Under the conditions of our experiments, taking them from their home tank and placing them in a behavioral chamber constitutes agitation, which has been shown to result from the release of serotonin into the bloodstream, an effect that lasts 15-20 minutes (Willard 1981). The stepping behavior on uniform surfaces of different grits (Fig. 2) diminishes with a similar time constant. The sensory input that produces the increased number of head withdrawals may also prolong the modulatory effects of serotonin or dopamine, a neuromodulator that activates crawling behavior in whole leeches and semi-intact leeches (Crisp et al. 2012), and activates the crawling pattern generator in isolated segmental ganglia (Puhl and Mesce 2008).

#### Comparisons with other species

The head movements that *H. austinensis* uses to determine a favorable substrate surface is similar to the “scanning” movements used by medicinal leeches (*Hirudo verbana*) to locate their prey (Harley and Wagenaar 2014). Many other animals use body extensions such as whiskers and antennae to probe their near-field environment using mechanosensors to determine whether they are about to run into an object (Harley and Ritzmann 2009), to cross over a hole in the ground (Blaesing and Cruse 2004), or even to determine what the object is (rat whisking, Bush et al., 2016). In fact, in a darkened room—especially an unfamiliar one--people use the same sorts of scanning movements with their arms and legs to avoid running into walls, doors, and furniture.

At the level of touch sensation, humans, in describing the texture of something that they touch with their finger tips, usually mention its texture—how rough or smooth--the surface feels (Tiest 2010). People can discriminate quite fine differences between different coarseness. They can tell the difference between 600 and 1500 grit sandpapers, for instance, that differ in particle size by an average of just 10 microns (16 vs 6 microns), but only if they move their fingertips over the surface. Interestingly, for coarse sandpapers (120 vs 80 grit, which have, on average, particles of 116 vs 192 microns), people do nearly as well using static touch as they do by brushing their fingers over the surface (Hollins and Risner 2000). Similar psychophysical measurements have been made on experimental animals, using finely controlled movements of precisely machined surface textures (Eck et al. 2013). These types of studies have shown that the discriminations among smooth textures are likely made by the vibrations of the skin caused by the surfaces that move along the skin, thereby activating a particular class of tactile receptors (Pacinian corpuscles) that responds selectively to objects lightly moving over the surface of the skin (Hollins and Risner 2000; Grant et al. 2014). These responses to moving stimuli are characteristic of leech T cells (NIcholls and Baylor 1968; Lewis and Kristan 1998b; Pirschel and Kretzberg 2016), so it will be interesting to determine whether these neurons are selectively activated by the scanning movements made by leeches as they scan their environment. Because we have found that T and P neurons in *Hirudo* and *Helobdella* express many different types of orthologous transcripts (Heath-Heckman et al., in preparation), it should be possible to use genome editing approaches to selectively modify the features of specific cell types and then observe the consequences of these manipulations on the behavior of the intact animal.

## ACKNOWLEDGEMENTS

We thank Lidia Szczupak and members of the Weisblat lab for helpful comments on the manuscript.

